# Use of Hypochlorous Acid in Infections: Bibliometric Analysis of Scientific Literature

**DOI:** 10.1101/2024.11.06.622311

**Authors:** Fatih M. Ateş

**Affiliations:** Bayburt Universitesi

**Keywords:** Hypochlorous acid, infection, use, bibliometric analysis, bibliometrics

## Abstract

**Background:** Historically, chemicals such as alcohols, aldehydes, chlorinated compounds and hypochlorous acid have been used to eliminate or inactivate infectious agents. Since bibliometrics is a powerful tool for uncovering scientific literature, we decided to conduct a bibliometric analysis of the literature on the use of hypochlorous acid in infections. Emerging trends and common patterns in research were explored, collaborations and networks were monitored and future directions were predicted.

**Methods:** The Scopus database was searched for documents published between 1982 and 2024 using the keywords “hypochlorous acid”, “infection” and “use”. Data analysis was performed using R-Studio 2024.09.0 Build 375 software with a machine learning bibliometric method based on the bibliometrix R package. The most relevant authors were measured by the number and fractional number of documents written. Author productivity was analyzed with Lotka’s law. Bradford’s law was applied to identify the core of journals focusing on the topic under consideration. The field of mainstream themes included isolated topics (niche themes), new topics (emerging themes), hot topics (engine themes) and core topics (core themes).

**Results:** A total of 574 documents were found, the vast majority of which were original articles (74%). The annual growth rate was 5.08%. Overall, 424 journals published more than one document. Most of the corresponding authors work in the United States. Neutrophils, animals and hydrogen peroxide are emerging themes, while hypochlorous acid, human, article represent the basic themes.

**Conclusions/Applications for Practice:** The interests of infection specialists have diversified over time and metabolism, reactive oxygen metabolites, anti-infective agents have been added.

## Background

From past to present, human beings have had to struggle with countless infectious agents. Sterilization and disinfection processes have been used to eliminate or neutralize these infectious agents. Sterilization refers to the complete elimination of microbiological agents, including spores, from inanimate surfaces by physical or chemical methods, while disinfection refers only to the elimination of infectious agents. In disinfection processes, factors such as pre-cleaning of the application area, type and intensity of infection, pH, temperature, disinfectant concentration, application time, etc. are important factors that ensure the success of disinfection (Ateş, 2020). The destruction of pathogenic microorganisms in living tissues with chemicals is called antisepsis. While heat, radiation, filtration, gas vapor and chemicals can be used for sterilization; physical methods can be used for pasteurization. Alcohols, aldehydes, chlorinated compounds and oxidizers can be used for chemical disinfection. In addition to these, chemicals such as chlorine, hypochlorous acid and sodium dichloroisocyanurate (NaDCC) can also be used for disinfection (Külekçi, 2005). Regulation of environmental factors is not enough to protect against infections; strengthening the immune system, eating a balanced diet, paying attention to personal hygiene and taking precautions during travel are also effective protection methods (Ateş, 2020).

It is usual to prefer chemicals to be used in disinfection that do not cause corrosion on the surface they come into contact with, are non-toxic and low priced. Hypochlorous acid (HOCl) is widely used as an economical and effective disinfectant against pathogenic microorganisms in all mammals since it can be produced by electrolysis of water and NaCl. Neutrophils, eosinophils, B-lymphocytes produce HOCl in response to infections with nicotinamide adenine dinucleotide phosphate oxidase bound to the mitochondrial membrane (Kettle & Winterbourn, 1997). The resulting HOCl reaches maximum antimicrobial activity at pH=3-6 (Wang et al., 2007). In aqueous solution, HOCl decomposes into H+ and OCl^−^ and denatures proteins. HOCl renders viruses harmless by forming chloramine and nitrogen radicals and breaking double-stranded DNA (Winter et al., 2008). It can be stated that the destruction of infectious agents in humans and the ecosystem they live in through HOCl is a safe method. The use of HOCl in the prevention of infections will enable national and international collaborations, networking capabilities and organizational activities to be carried out. In this study, a bibliometric review was conducted to analyze the literature, investigate trends, patterns in collaborative research, track collaboration networks and predict future research directions between 1982 and 2024 on the use of HOCl in infections.

## Methods

Today, various databases such as Web of Science, Scopus (WoS), Google Scholar, Microsoft Academic, Crossref, Dimensions and CiteSeer can be used for bibliometric analysis. Google Scholar has been compared to other databases such as WoS and Scopus by a number of researchers because it is the bibliography database most preferred by researchers. These researchers found significant overlap in terms of content, number of articles, typology, number of conferences and proceedings (Aguillo, 2012; Bar-Ilan, 2010; García-Pérez, 2010; Jacsò, 2010; Li et al., 2010). There are several reports arguing that Scopus is a strong competitor of WoS in bibliometrics. Similarly, compared to WoS, Scopus is cited as having a broader coverage and possibly a larger research area, especially in the social sciences (Li et al., 2010; Mongeon & Paul-Hus, 2016). Bass J et al. stated that Scopus is an important source for curated, high quality, bibliometric data analysis in academic research (Baas et al., 2020). Therefore, in our study, the Scopus database was used for the analysis. A comprehensive search was conducted in the Scopus database between July 1982 and September 2024. Search fields included article title, abstract and keywords. The keywords “Hypochlorous Acid”, “Infection”, “Use” were used in the database searches. Bibliographic metadata were downloaded in BibTex format and exported in Excel format in R environment (R-Studio 2024.09.0 Build 375 software). Bibliometric analysis of the excel file obtained from the Scopus database was performed through the R-Studio program.

A bibliographic data frame was created and the selected documents contain bibliographic attributes such as authors’ names, links, title, keywords, journal, year, volume, number, page, editor(s) and number of citations. The “summary ()” function was used to summarize the main results. Data collection covered basic information (annual scientific production, annual average citations), sources (most relevant sources, most cited sources, source dynamics), authors (most relevant authors and their links, country of the relevant author, country-specific production) and documents (most globally cited documents, most frequently used keywords, word dynamics). Annual growth rate was used to describe the rate of progress of scientific production over time.

The most relevant authors were analyzed both in terms of number and contribution in co-authorship. Author productivity was analyzed using Lotka’s law. Bradford’s law was applied to identify the core of journals focusing on the topic under consideration. Multi-country production (MCP) indicates the number of documents where at least one co-author is linked to a country other than the first author. The most relevant links (frequency distribution of all co-authors’ links for each document) were based on link elements disambiguated by applying semantic similarity. The country of the corresponding author was used to list the most relevant countries according to the corresponding author. Country scientific production refers to authors appearing by country affiliation (number of documents indicates authors appearing by country affiliation). The most productive country was determined by the institution to which the first author was affiliated. A correspondence analysis and clustering approach was used to identify common shared keywords and author/institution relationships across research themes. A thematic map was drawn by applying a clustering algorithm on the keyword network. The mainstream themes area includes isolated topics (niche themes), new topics (emerging themes), hot topics (engine themes) and core topics (core themes). Each bubble represents a network cluster. Keywords with the highest occurrence value were used to define the bubble name. Bubble size was proportional to cluster word occurrence and its position was adjusted according to cluster centrality and density. The function “Biblioshiny ()” was used to plot all results.

## Results

### Overview

We collected 574 documents, mostly articles (n = 424; 74%) and review articles (n = 97; 16%). A total of 2571 authors contributed to these documents from 1982 to 2024, with an average of 6 co-authors per publication. The annual growth rate was 5.08%, resulting in an increase in scientific production in the literature. While 4 documents were published in 1982, the number of these documents increased to 574 in 2024. The average annual total number of citations is between 2-3 citations. The highest average total citations per article was 11.24 citations in 2017, while 1985 was the lowest with 0.15 average citations.

### Sources

Overall, 424 journals published one or more documents. As shown in Figure 1, 30 journals published about 33% of all retrieved documents. “Journal of Hospital Infection” published the most articles with 23 publications. “Plos One” published 22 articles, ‘Infection and Immunity’ published 21 articles, and ‘Journal of Wound Care’ published 18 articles. Among the top 5 journals, there was an increase in publications on the subject in 2012-2024 compared to 1982-2012. The most cited journal on the subject was “Nutrients” with 1151 citations.

**Figure 1.**
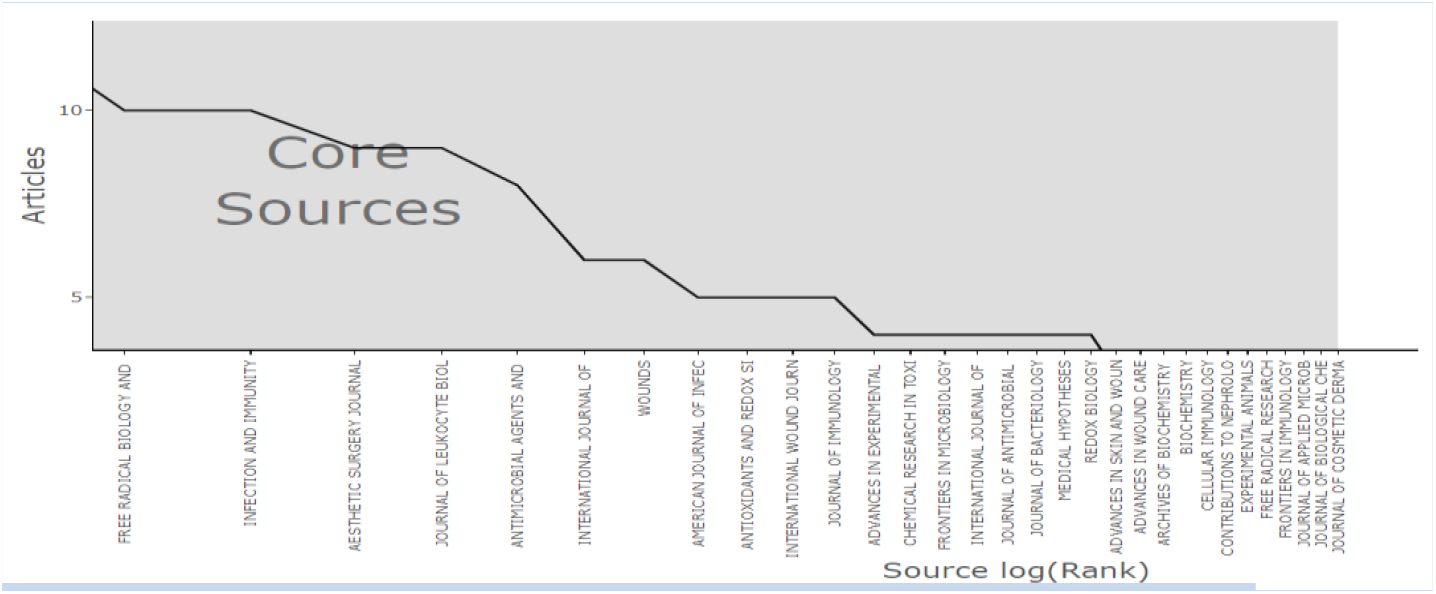
Source Clustering Through Bradford’s Law

### Authors, affiliations, countries

The most relevant authors were Kettle A.J., Beyenal H. and Patel R. with a fractional frequency of 16 (3.2), 13 (1.72) and 13 (1.72) documents, respectively. The frequency distribution of scientific productivity identified that there were several “core” authors (n = 287; 11.1%) who wrote at least two papers and “occasional” authors (n = 2284; 88.9%) who published only one paper. The countries of the top twenty authors are presented in Figure 2. According to the MCP ratio, India, Turkey, Ireland and Ireland show a low rate of international collaboration, while the USA, New Zealand, United Kingdom have the highest rate. Country-specific production is shown in Figure 2, with the USA being the most productive country with 542 documents. The most relevant institutions were Washington State University, University of Otago Christchurch, Louisiana State University Health Sciences Center.

**Figure 2.**
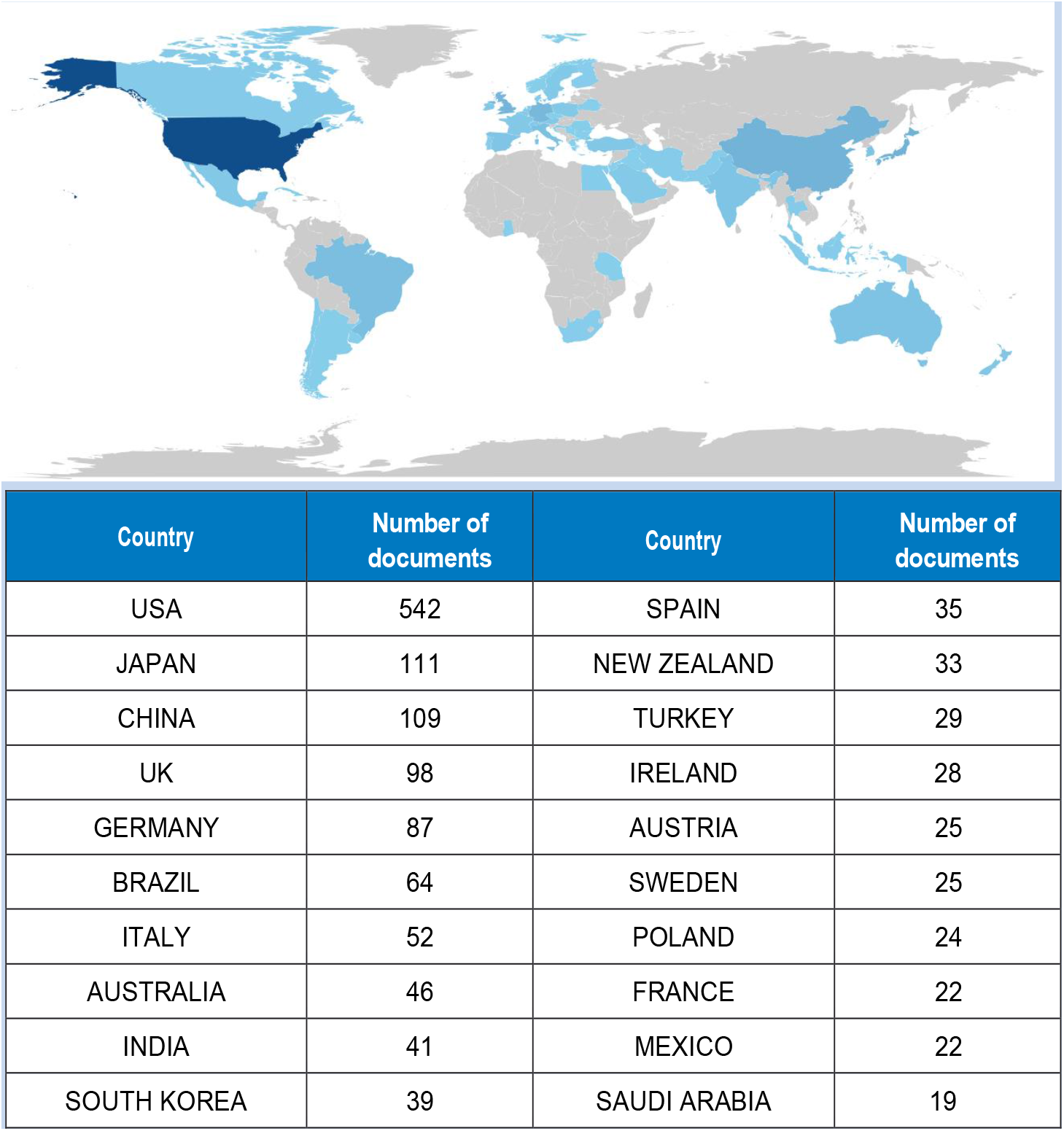
Country-Specific Production

### Documents

The top ten most cited papers are listed in Table 1 (Carr & Maggini, 2017; Aguillo, 2012; Bar-Ilan, 2010; García-Pérez, 2010; Jacsò, 2010; Li et al., 2010; Mongeon & Paul-Hus, 2016; Baas et al., 2020; Boyce, 2016; De Larco et al., 2004; El-Benna et al., 2008). The most cited documents in the primary search strategy (“Hypochlorous Acid” and “Infection”) were retained in the sensitivity analysis with the addition of the term “use”. The vast majority of these ten articles are review articles. The first ranked article is “Vitamin C and Immune Function” with 1151 global citations (Carr & Maggini, 2017).

**Table 1.** Top Ten Most Global Cited Documents

| Paper | DOI | Total citation | TC per year | Normalized TC |
| --- | --- | --- | --- | --- |
| CARR AC, 2017, NUTRIENTS | https://doi.org/10.3390/nu9111211 | 1151 | 143,875 | 12,802 |
| EVANS P, 2001, BR J NUTR | https://doi.org/10.1079/bjn2000296 | 435 | 18,125 | 2,691 |
| KNIGHT JA, 2000, ANN CLIN LAB SCI |  | 425 | 17,000 | 2,459 |
| JAESCHKE H, 2006, AM J PHYSIOL GASTROINTEST LIVER PHYSIOL | https://doi.org/10.1152/ajpgi.00568.2005 | 424 | 22,315 | 3,072 |
| SCHULLER-LEVIS GB, 2003, FEMS MICROBIOL LETT | [https://doi.org/10.1016/S0378-1097(03)00611-6](https://doi.org/10.1016/S0378-1097%2803%2900611-6) | 368 | 16,727 | 2,947 |
| DE LARCO JE, 2004, CLIN CANCER RES | https://doi.org/10.1158/1078-0432.CCR-03-0760 | 359 | 17,095 | 3,247 |
| WINTERBOURN CC, 2013, ANTIOXID REDOX SIGNAL | https://doi.org/10.1089/ars.2012.4827 | 355 | 29,583 | 6,145 |
| BARKER J, 2004, J HOSP INFECT | https://doi.org/10.1016/j.jhin.2004.04.021 | 325 | 15,476 | 2,939 |
| EL-BENNA J, 2008, SEMINIMMUNOPATHOL | https://doi.org/10.1007/s00281-008-0118-3 | 293 | 17,235 | 4,060 |
| BOYCE JM, 2016, ANTIMICROB RESIST INFECT CONTROL | https://doi.org/10.1186/s13756-016-0111-x | 292 | 32,444 | 5,620 |

Note. DOI: Digital object identifier; TC: total citations

Most of these articles (n = 10) were published between 2000 and 20117, while more recent articles (2018-2024) were limited in number, possibly due to the shorter citation timeframe. After removing the search strategy terms “hypochlorous acid” and “infection”, keyword analysis revealed that “human”, “article” and “humans” were the three most frequently used words, occurring 436, 335 and 324 times, respectively. The word cloud is shown in Figure 3.

**Figure 3.**
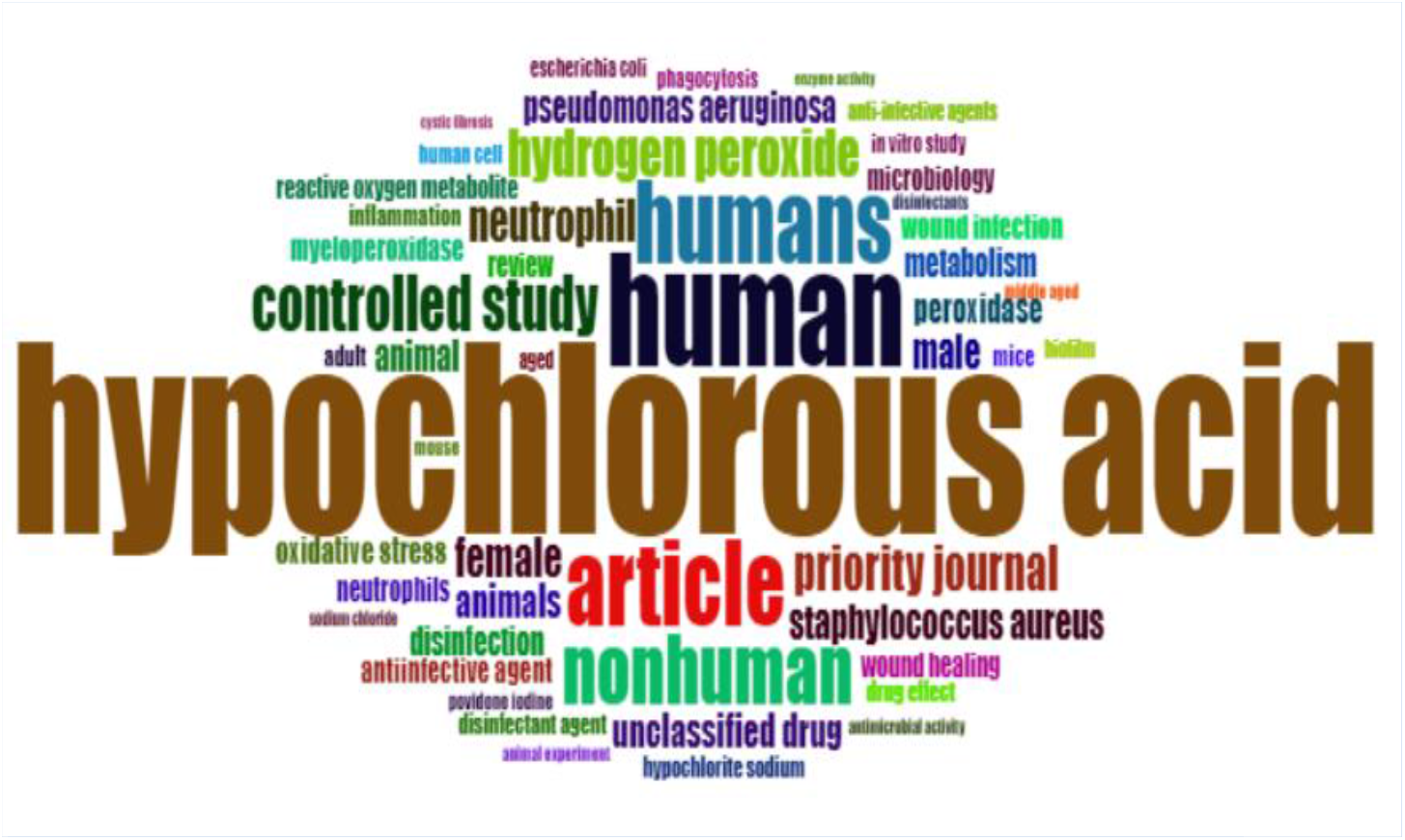
Word Cloud

### Conceptual structure

The main themes and trends are shown in Figure 4 and Figure 5. The thematic map (Figure 4) shows that neutrophils, animals and hydrogen peroxide are emerging themes, while hypochlorous acid, human, article represent the main themes. Figure 4 shows the evolution of the main thematic areas and their relationships over four different time periods: 1982-1994, 1995-2004, 2005-2014 and 2015-2024. Over time, the ‘human’ theme links have varied from hypochlorous acid, disinfection, microbiology, neutrophil, anti-infective agent. ‘Human’ and ‘article’ merged under ‘hypochlorous acid’ in 1995-2004. ‘Taurine’ started as a niche theme and merged into ‘hypochlorous acid’ in 2005-2014. ‘Anti-infective agent’ first emerged in 2005-2014, and continued to attract attention in the following decade. The country collaboration network (Figure 6) shows country collaboration based on publications. Line thickness indicates closeness of collaboration. The US has a strong connection with Japan and the United Kingdom (UK). The tendency of European countries to cooperate with each other is evident.

**Figure 4.**
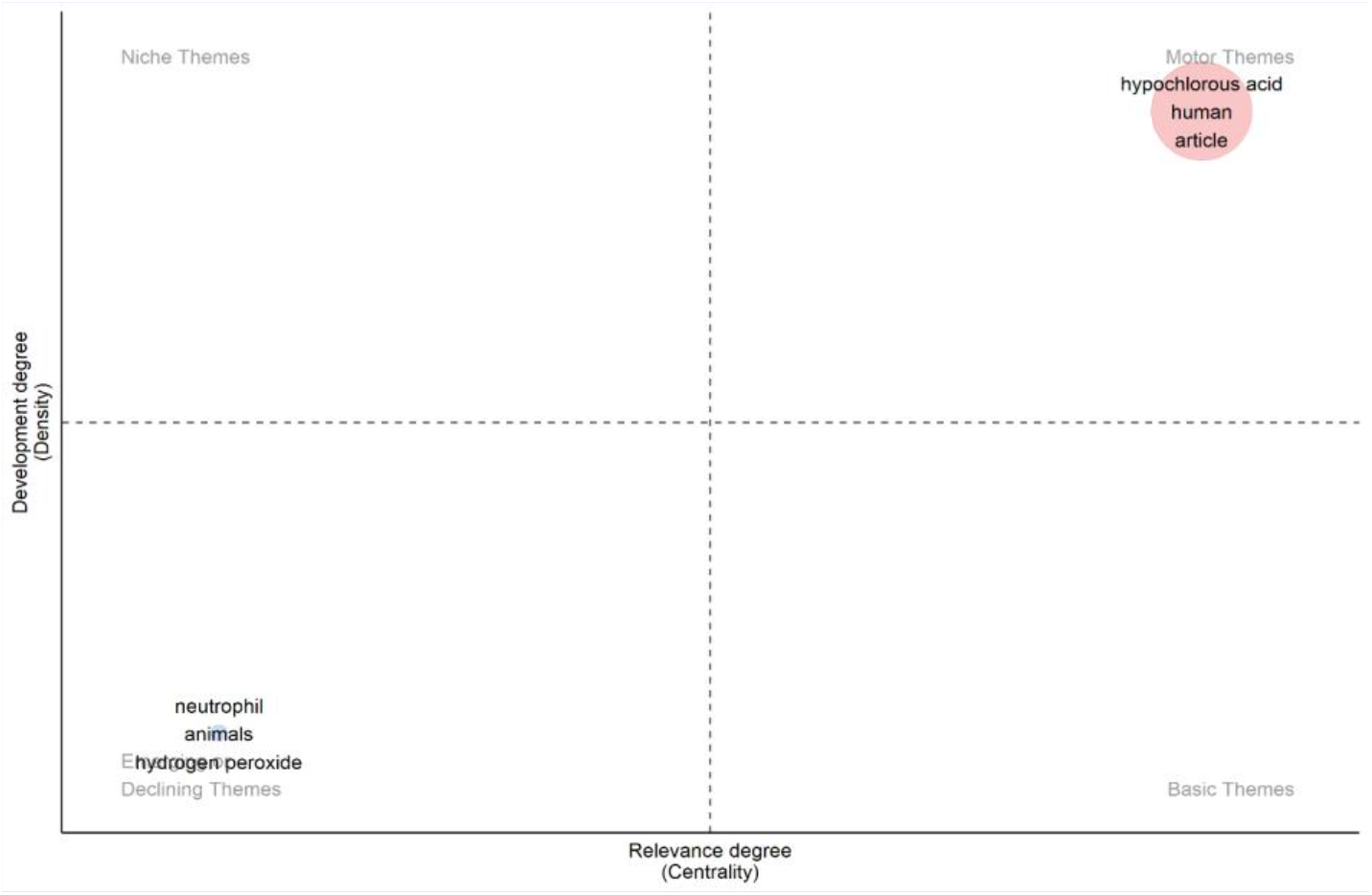
Thematic Map

**Figure 5.**
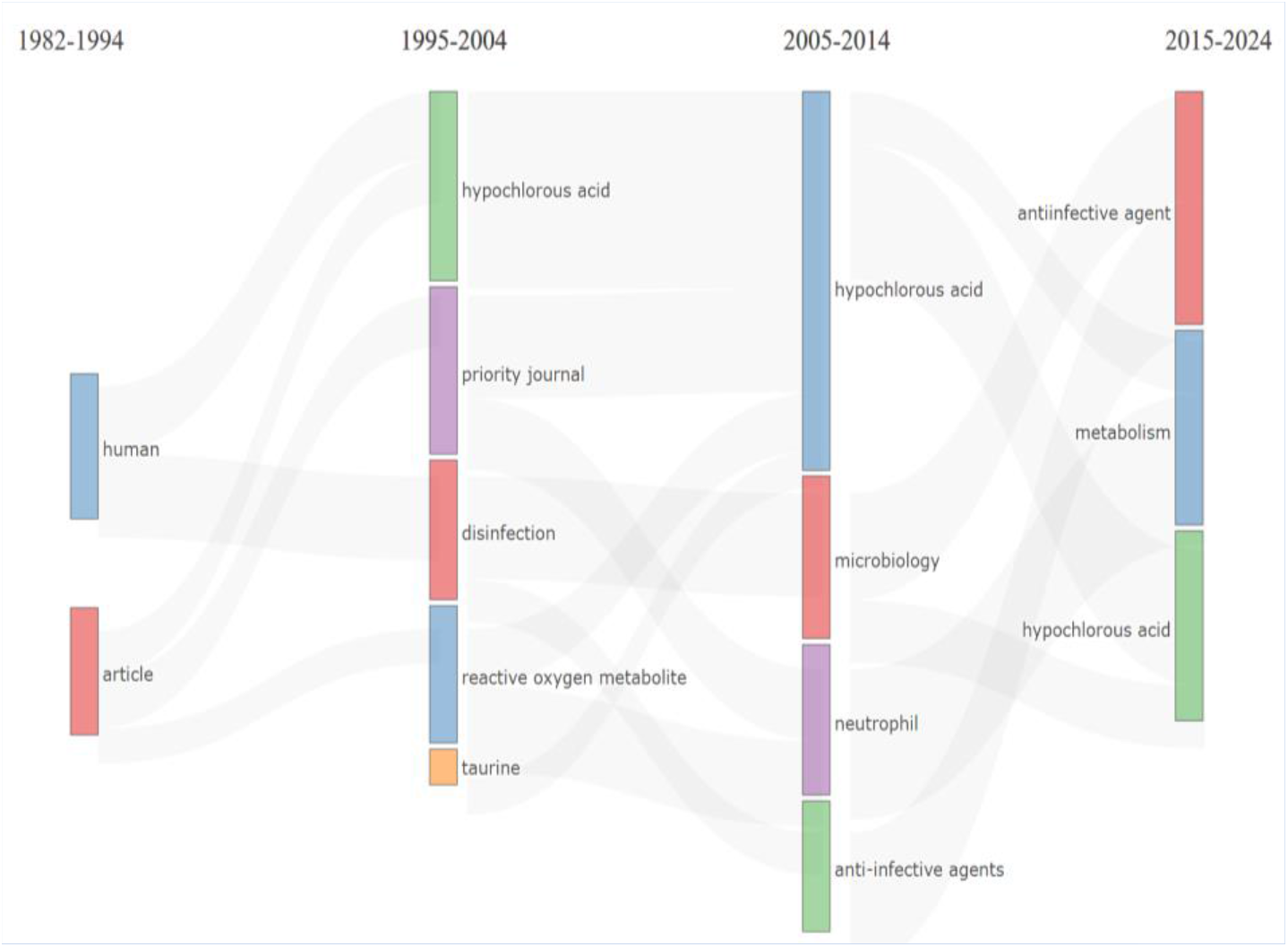
Thematic Evolution

**Figure 6.**
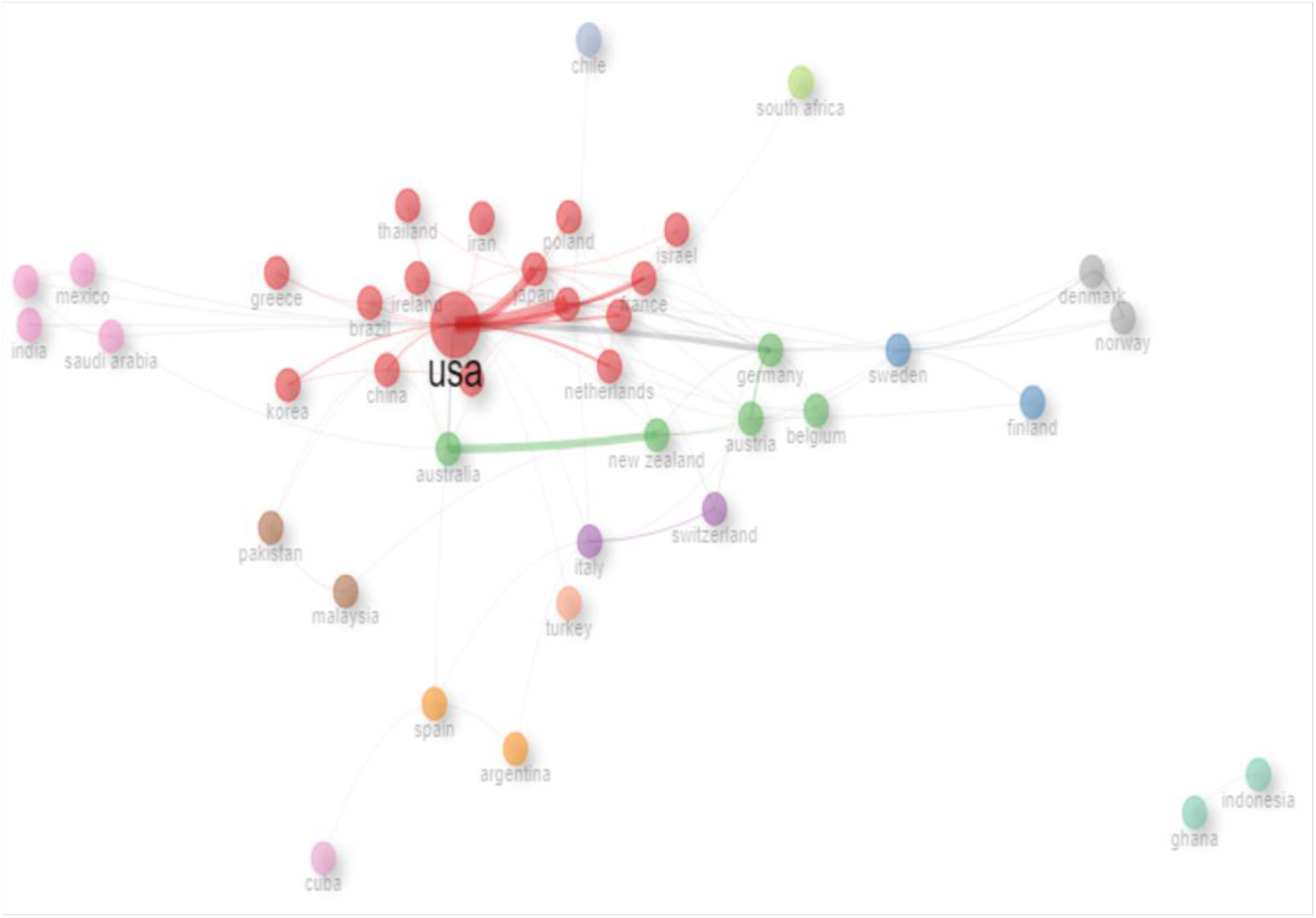
Collaboration Network

## Discussion

A bibliometric analysis of the scientific literature on the use of HOCl in infections between 1982 and 2024 revealed an average annual increase in scientific production of approximately 6%, with 32 articles published up to October 2024. Of the 574 documents published between 1982 and 2024, 74% are original articles, underlining the novelty of the specific scientific production. About one fifth of all published documents are published by a limited number of highly ranked journals with an impact factor above 2, covering the fields of infection, microbiology and immunology, along with HOCl. Journals dedicated to infections showed a sharp increase in the number of publications on HOCl between 2012 and 2024, indicating a higher scientific interest in neutrophils as part of the multidisciplinary management of infections.

The geographical distribution of scientific publications spans all continents and reflects the high incidence of infection worldwide. The dominant centers for scientific production are North America (USA, Canada and Mexico), Europe (Germany, Italy, UK, Spain) and Asia (China, India, South Korea) and Europe. The USA, Canada, the UK and China have both high quantitative scientific output and the capacity to form an effective collaborative network that is not dispersed but rather selected. Continental European countries can sustain high scientific productivity and tend to implement scientific cooperation at the European level. Scandinavian countries have their own unique collaborative networks. Other countries (Turkey, India, Korea) are able to sustain high scientific productivity through a diffuse ‘go it alone’ approach based on national collaborations. The geographical distribution of scientific production shows that there are productive academic institutions in North America (Washington State University, Louisiana State University Health Sciences Center, University of Iowa), Europe (Freie Universität Berlin, University of Copenhagen) and New Zealand (University of Otago Christchurchen). A limitation of the present analysis is that collaboration is measured using a simple proxy of co-authorship, which may not reflect an active scientific network or the scientific value of the published work. Furthermore, the search strategy we used was chosen as a balance between comprehensiveness and usability to limit the background noise potentially introduced by the addition of other more comprehensive keywords (such as infection or use).

Some of the most cited publications between 2000 and 2024 are directly focused on infection. Articles by Carr, Evans, Knight et al (Vitamin C and Immune Function, Micronutrients: Oxidant/Antioxidant Status and Review: Free Radicals, Antioxidants, and the Immune System) addressed the use of hypochlorous acid in infections (Carr & Maggini, 2017; Evans & Halliwell, 2001; Knight, 2000). Similarly, Jaeschke ‘Mechanisms of Liver Injury. II. Mechanisms of neutrophil-induced liver cell injury during hepatic ischemia-reperfusion and other acute inflammatory conditions’ showed that it is possible to prevent infections by HOCl production in neutrophils (Jaeschke, 2005). Winterbourn & Kettle ‘Redox Reactions and Microbial Killing in The Neutrophil Phagosome. Antioxidants & Redox Signaling’ found, in line with previous researchers, that neutrophils rapidly kill most ingested microorganisms by a myeloproxidase-dependent mechanism that is almost certainly HOCl-dependent (Winterbourn & Kettle, 2013). Boyce, in his study ‘Modern Technologies for Improving Cleaning and Disinfection of Environmental Surfaces in Hospitals’, stated that hydrogen peroxide-based liquid surface disinfectants, a combination product containing peracetic acid and hydrogen peroxide, are effective alternatives to currently widely used disinfectants and that electrolyzed water (hypochlorous acid), cold atmospheric pressure plasma have potential for use in hospitals (Boyce, 2016). HOCl is used in general cleaning and disinfection processes in the agricultural sector, food sector, medical sector (Park et al., 2007; Stroman et al., 2017). If HOCl is applied by fogging method, it is antibacterial and antiviral (Zhao et al., 2014). Due to the fact that HCOl is cheaper than other disinfectants and has a wide range of antimicrobial effects, different, especially medical uses have increased and diversified. One of the most important reasons for its widespread use is that HOCl can also be produced by the human defense system. Therefore, it can be said to be more natural when other cleaning and antimicrobial agents are considered (Ateş, 2020). The major limitation of the study is linked to its very nature. Bibliometric analysis provides quantitative information with very high or low predictive potential. The quality of the data may be biased due to the lack of standardization of the elements investigated, such as their links. The use of only documents from the Scopus database is insufficient to present the current work on infection and all global research activities related to HOCl. In addition, quantitative data, especially the number of citations, can be affected by the number of years for which publications can collect references. Furthermore, many studies may report updates and multiple publications, different endpoints and different follow-up times.

The research objectives for the use of HOCl have changed over time to reflect the worldwide evolution of the fight against infections. The management of infection is now based on a multidisciplinary integration of therapies, including systemic and targeted agents and surgery. The primary interests of infection specialists have evolved over time, adding safety, health-related quality of life, sustainability of therapies and combination to systemic therapy to overall efficacy and effectiveness and treatment outcomes. The data presented here provides a unique window to appreciate the eclectic nature of HOCl as a discipline in the prevention of infections, highlighting the fact that technology and clinical management cannot evolve without each other in a process of continuous growth and optimization.

